# Structure-Aware Annotation of Leucine-rich Repeat Domains

**DOI:** 10.1101/2023.10.27.562987

**Authors:** Boyan Xu, Alois Cerbu, Daven Lim, Christopher J Tralie, Ksenia Krasileva

**Affiliations:** Center for Computational Biology, University of California Berkeley, Berkeley, CA 94720, U.S.A; Department of Mathematics, University of California Berkeley, Berkeley, CA 94720, U.S.A; Department of Plant and Microbial Biology, University of California Berkeley, Berkeley, CA 94720, U.S.A; Department of Mathematics and Computer Science, Ursinus College, Collegeville, PA, USA

## Abstract

Protein domain annotation is typically done by predictive models such as HMMs trained on sequence motifs. However, sequence-based annotation methods are prone to error, particularly in calling domain boundaries and motifs within them. These methods are limited by a lack of structural information accessible to the model. With the advent of deep learning-based protein structure prediction, existing sequenced-based domain annotation methods can be improved by taking into account the geometry of protein structures. We develop dimensionality reduction methods to annotate repeat units of the Leucine Rich Repeat solenoid domain. The methods are able to correct mistakes made by existing machine learning-based annotation tools and enable the automated detection of hairpin loops and structural anomalies in the solenoid. The methods are applied to 127 predicted structures of LRR-containing intracellular innate immune proteins in the model plant *Arabidopsis thaliana* and validated against a benchmark dataset of 172 manually-annotated LRR domains.

**Author summary:** In immune receptors across various organisms, repeating protein structures play a crucial role in recognizing and responding to pathogen threats. These structures resemble the coils of a slinky toy, allowing these receptors to adapt and change over time. One particularly vital but challenging structure to study is the Leucine Rich Repeat (LRR). Traditional methods that rely just on analyzing the sequence of these proteins can miss subtle changes due to rapid evolution. With the introduction of protein structure prediction tools like AlphaFold 2, annotation methods can study the coarser geometric properties of the structure. In this study, we visualize LRR proteins in three dimensions and use a mathematical approach to ‘flatten’ them into two dimensions, so that the coils form circles. We then used a mathematical concept called winding number to determine the number of repeats and where they are in a protein sequence. This process helps reveal their repeating patterns with enhanced clarity. When we applied this method to immune receptors from a model plant organism, we found that our approach could accurately identify coiling patterns. Furthermore, we detected errors made by previous methods and highlighted unique structural variations. Our research offers a fresh perspective on understanding immune receptors, potentially influencing studies on their evolution and function.

## Introduction

Solenoid domains are a class of protein structures defined by a repeating helical arrangement of their backbone chain. These domains are found in a diverse range of proteins and play important roles in a variety of biological processes, including protein-protein interactions, molecular recognition, and scaffolding [1]. The coil shape of solenoid domains arises from a repeating motif of amino acid residues, known as *tandem repeat units*. The specific amino acid sequence and length of the repeating unit can vary between solenoid domains, resulting in differences in the overall structure and function of the domain. The modular nature of solenoid domains allows for the construction of complex structures by combining different domains in a predictable and controlled manner [2].

Leucine-rich repeat (LRR) domains are a type of curved solenoid domain with repeated units of about 20 - 30 residues long which contain leucine residues in a beta-strand conformation. These domains are found in a wide range of proteins, including cell surface receptors, enzymes, and structural proteins, and are known to play important roles in protein-protein interactions, signal transduction, and immune recognition [4].

Leucine-rich repeats play a critical role in the function of the NOD-like receptor (NLR) family of proteins in the innate immune system of plants and animals [5]. NLRs are intracellular immune receptors that recognize pathogen-derived molecules and activate downstream signaling pathways to initiate an immune response. NLRs are involved in the recognition of a wide range of pathogens, including bacteria, fungi, and viruses. NLRs typically consist of three domains: an N-terminal domain, a central nucleotide-binding domain, and a C-terminal LRR domain. The LRR domain is responsible for recognizing and binding to pathogen-derived molecules, such as effector proteins or pathogen-associated molecular patterns (PAMPs) [6]. In particular, the LRR domains of plant NLRs are highly diverse and can recognize a wide range of pathogen-derived molecules, allowing plants to mount a robust and specific immune response to a broad range of pathogens. Understanding LRR domains in plant NLRs is important for developing strategies to enhance plant immunity and improve crop resistance to pathogens.

The concave surface of the leucine-rich repeat domain is generally responsible for binding to ligands [7]. The amino acid residues on the concave surface of the LRR domain form a specific pattern of hydrophobic, polar, and charged residues that can interact with specific ligands, such as proteins, peptides, carbohydrates, or nucleic acids. The specificity of ligand binding by LRR domains is determined by the overall shape and chemical properties of the concave surface, which can be highly variable between different LRR-containing proteins [8][9]. Additionally, LRR domains can contain variable regions and insertions that can modify the binding specificity and affinity of the domain. More recently, studies such as [10] have shown that “post-LRR” domains which lie at the C-terminal end of the LRR are required for successful plant immune response. Accurate annotation of these domains and their constituent repeat units is thus essential to understanding the components which govern protein shape and binding specificity.

Existing methods for annotating LRR domains give unreliable and inconsistent results due to irregularities in sequence motifs. Profile hidden Markov models (HMMs) are widely used, e.g. by HMMER [11], to annotate protein domains in genomic sequences, but they are sensitive to the size and diversity of the protein family being analyzed and do not perform accurately for rapidly-evolving, highly-divergent families such as LRR [12]. Profile HMMs are also unable to delineate tandem repeat units.

An existing tool, LRRPredictor [13], uses an ensemble of 8 machine learning classifiers to determine the residues which comprise the basic LRR motif of the form “LxxLxL” (where “L” refers to Leucine or other hydrophobic amino acid, and “x” can be any amino acid). We found that LRRPredictor often makes mistakes, particularly in identifying divergent motifs near the C- and N-terminal boundaries of the LRR. Because LRRPredictor, like an HMM, is trained on a specific set of LRR sequences taken from Protein Data Bank [14] (PDB), it incorrectly annotates LRR sequences which diverge from its training set.

With AlphaFold 2 [15], a deep-learning-based model, reliable protein structure prediction has become readily available, enabling domain annotation methods with direct access to geometric data from the protein. We leverage this geometric information to annotate essential features of the LRR domain: start/end position, post-LRR detection, repeat unit delineation, and structural irregularities.

From the perspective of differential geometry, a coiling curve in 3D space is characterized by a linearly increasing winding number around a core curve. We therefore detect the coiling LRR region, as the loci where the winding number is sufficiently close to a line of a fixed slope; the post-LRR domain is then decided as C-terminal sequence downstream from the point at which steady winding terminates. The methods section below describes our procedure for computing the winding number across the length of the protein. In contrast to HMM-based or other data driven techniques, our method is completely unsupervised and driven by simple mathematical methods.

## Methods

### LRR domain annotation via winding number calculation on 2D projection of normal bundle embedding

161 NLR protein sequences, i.e. *NLRome*, were obtained from *A. thaliana* Col-0 accession reference proteome. Of these 161 NLRs, 127 had Alphafold-predicted structures available on AlphafoldDB [15]. For each structure we extracted amino acid *α*-carbon positions to obtain a 3D backbone space curve.

To extract the LRR region from the rest of the protein, we leverage the solenoid geometry of the LRR domain in contrast to the post-LRR structure. The non-LRR regions do not wind regularly as the LRR does, so we annotate the LRR by computing the winding number of the coil around the core of the solenoid. We obtain the core curve of the coil by applying a Gaussian filter to the backbone 3D space curve. We then compute the tangent vector field of the core curve via a Gaussian derivative, i.e. applying a derivative Gaussian filter by convolving each component *γ*(*t*) of the core curve with the derivative

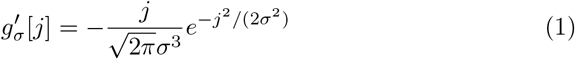

of a Gaussian via the formula 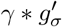 (in our case *σ* = 1). The normal bundle is obtained as the orthogonal complement to the tangent vector field. The backbone curve is embedded into the normal bundle which is then projected onto the Euclidean plane ℝ^2^ by parallel transport of an orthonormal frame along the core curve. Our algorithm for orthonormal framing is as follows:

#### Parallel transport algorithm for orthonormal framing

1. Randomly initialize 0^th^ pair of orthonormal 3D vectors, perpendicular to 0^th^ tangent vector on core curve. This is the current orthonormal frame.
2. To obtain the next orthonormal frame, project the current orthonormal frame onto the next normal plane. The resulting projection is likely not orthogonal; we replace it with the closest pair of orthonormal vectors. This can be done by computing the singular value decomposition of the 2 *×* 3 matrix and replacing its singular values with 1’s (the standard solution to the “Orthogonal Procrustes Problem” [16, 17]).
3. Repeat step 2 along the length of the core curve.

The parallel transport algorithm enables the identification of normal bundle fibers and thus projection onto a single 2D plane as shown in Figure 2. The winding number *w*(*s*) up to residue *s* is then computed using the formula:

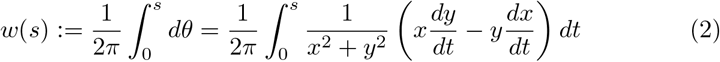

where *x*(*t*) and *y*(*t*) are the coordinates of the normal bundle projection as *t* ranges across the residues of the NLR protein sequence. We compute a discrete version of formula 2 by approximating 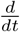 with a Gaussian derivative and integral with cumulative sum.

**Figure 1:**
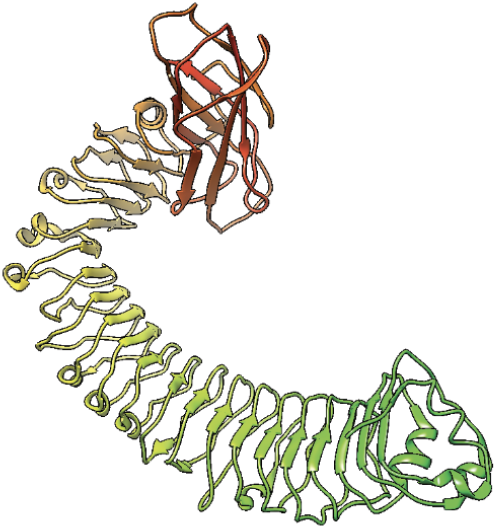
Cryo-EM structure of LRR from *A. thaliana* NLR RPP1 [3].

**Figure 2:**
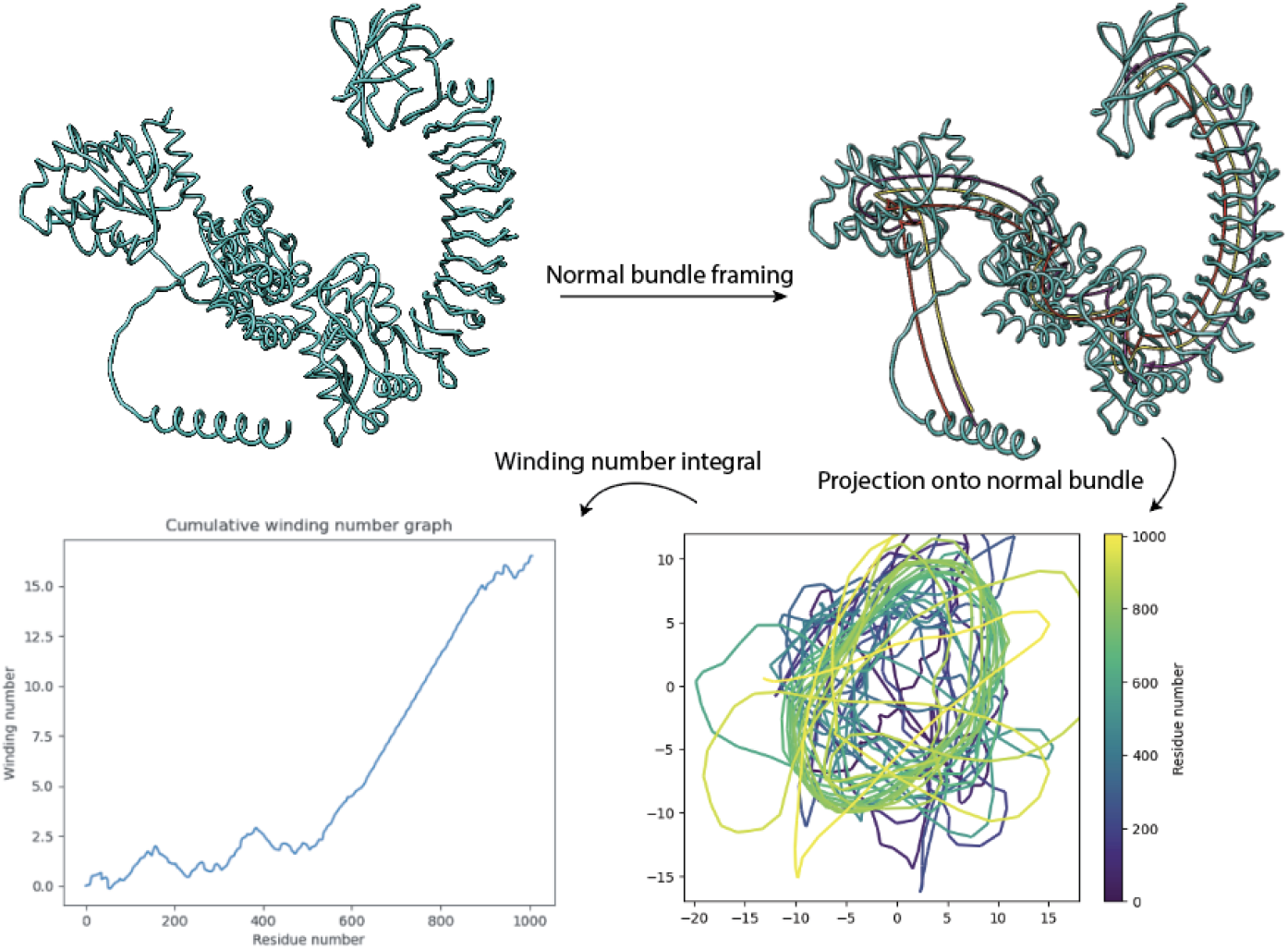
Embedding of protein backbone curve into normal bundle followed by projection onto an orthonormal frame yields a 2D curve containing a flattened slinky shown in lower right. The cumulative winding number, computed using the classic formula from calculus, is computed from the projection. Sloped linear segments of the winding number curve indicate coiling.

On the resulting cumulative winding number function we perform a least-squares piecewise linear regression with three pieces: the first being the best horizontal line, the second a sloped line of best fit, and third a horizontal line. The regressed function is similar to a ReLU with threshold or “clipped ReLU,” except that it need not be continuous at breakpoints. The horizontal lines flanking the sloped line represent regions of the NLR which are not LRR and do not have gradually increasing winding number. Therefore, the breakpoints in the regression estimate the start and end coordinates of the LRR domain. A plot of the winding number graph and regression, along with a comparison with HMM-based domain annotation from InterPro, is shown in Figure 3.

**Figure 3:**
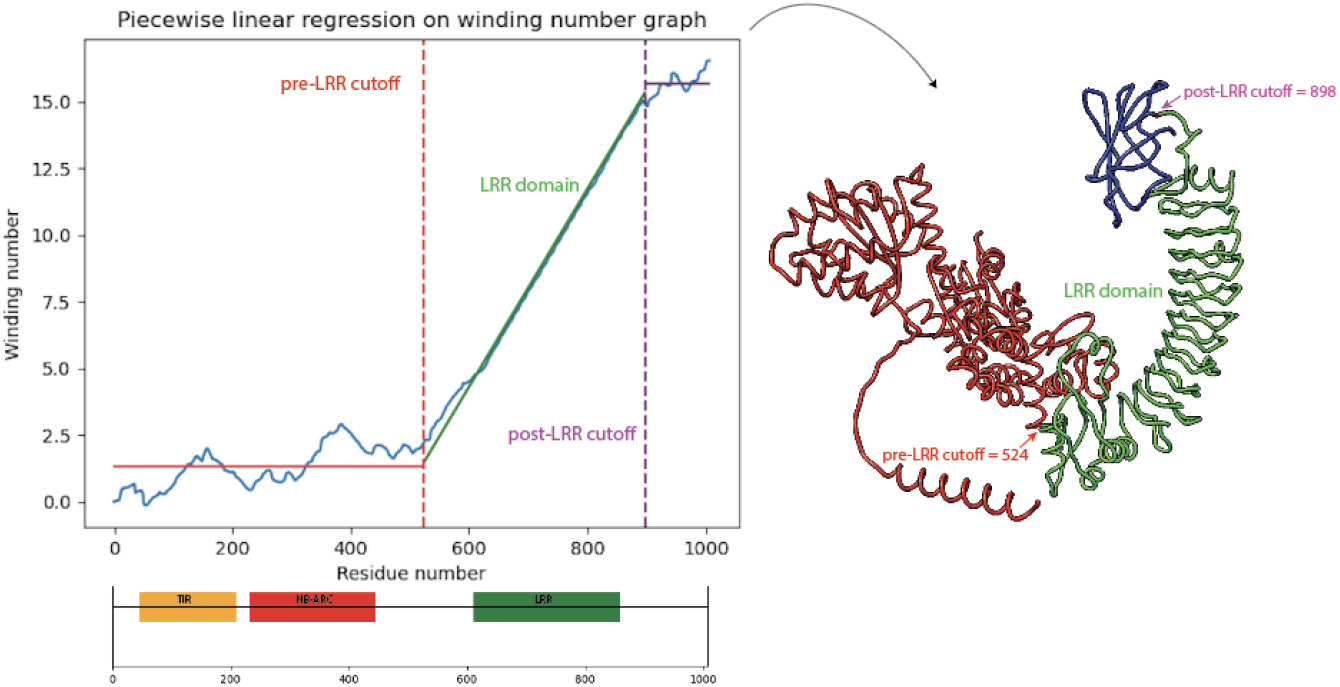
A discontinuous clipped ReLU function is regressed on the graph of the winding number function for *A. thaliana* NLR with TAIR [18] ID AT3G44400.1. The breakpoints of the regression yields the start and end positions of the LRR domain, highlighted in green. InterPro [19] domain annotations are shown below regression plot.

### Four-breakpoint regression detection of hairpin loops or mis-folding in LRR domain

A small number of proteins contain hairpin loops or other structural anomalies in the LRR domain which interfere with the clipped ReLU regression. Such proteins have a residual vector with high standard deviation in the sloped section. For these proteins, we run a similar piecewise linear regression with four break-points instead of two, registering a short noncoiling region along the otherwise solenoidal domain, represented by a short horizontal line in the piecewise-linear regression. This short horizontal line represents a large hairpin, insertion, or mis-folding within the LRR domain. See Figure 4.

**Figure 4:**
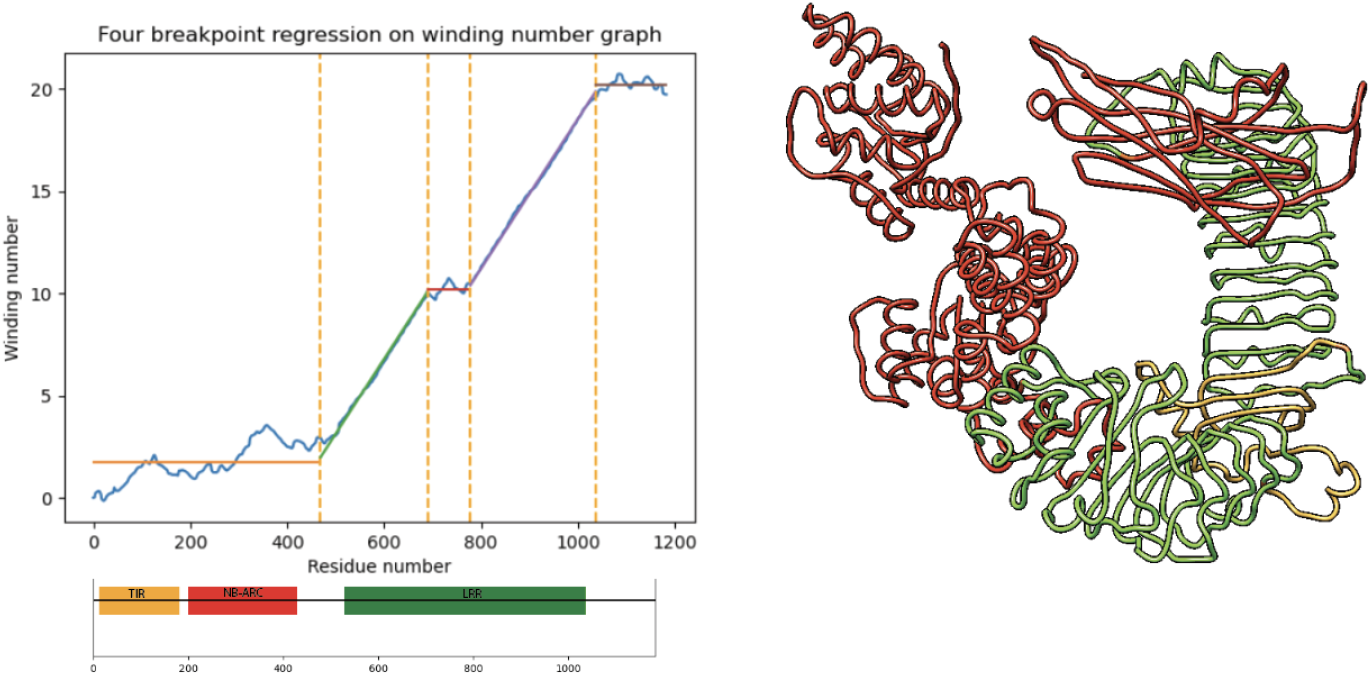
Four breakpoint piecewise linear regression enables detection of a non-coiling structure (highlighted in yellow at right) which deviates from the usual coiling in the LRR domain. Below regression plot, HMM-based InterPro domain annotations fail to detect non-coiling region within LRR domain. TAIR ID is AT1G72840.2.

### Eigenvectors of graph Laplacian on mutual nearest neighbors yield solenoid phase estimation

In the previous sections, we used piecewise linear regression on the cumulative winding number to isolate the LRR domain. In this section, we seek to obtain an angular coordinate, or *phase estimation*, on the LRR domain sequence. We first compute smoothened tangent vectors on the truncated LRR solenoid curve by convolving the position of the curve with the derivative of a Gaussian to obtain a tangent vector field. To accentuate periodic features in the coil [20], we perform a sliding window embedding of window size 24 (roughly the length of the LRR period) with delay time 1 on the tangent vector field, using the formula

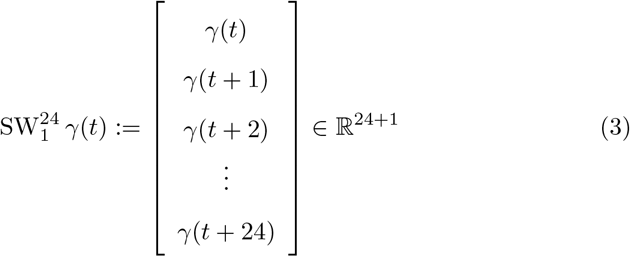

applied to each of the 3 components of the tangent vector field, resulting in a curve in 75-dimensional Euclidean space. We then construct a 50-mutual-nearest-neighbors graph on the sliding window embedding. From the mutual-NN graph we compute leading eigenvectors of the unweighted graph Laplacian, shown in Figure 5.

**Figure 5:**
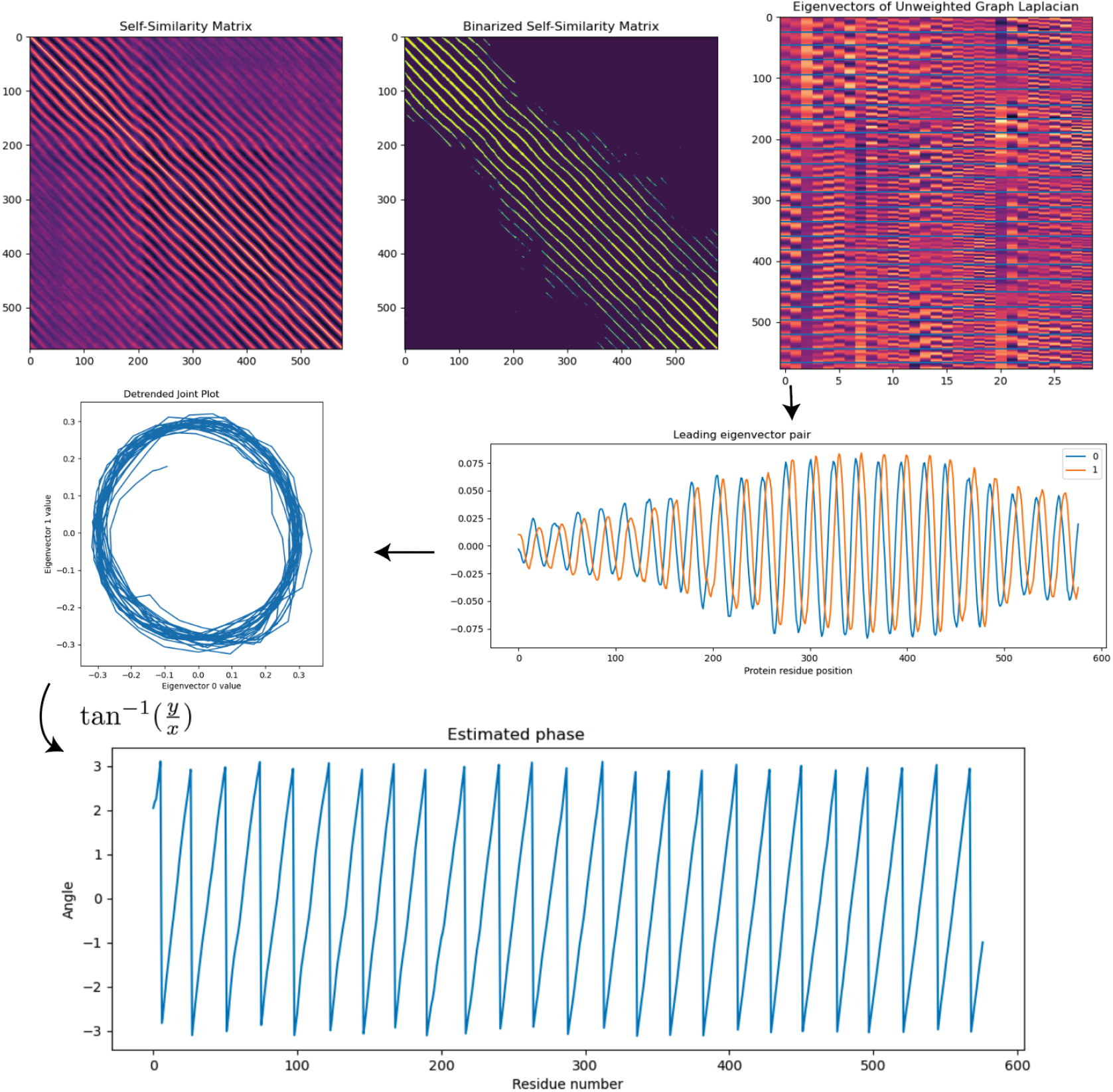
Graph Laplacian eigenvectors of mutual nearest neighbor graph on LRR solenoid curve tangent vectors. LRRPredictor residues are shown as blue horizontal lines on eigenmatrix plot. The 0^th^ and 1^st^ eigenvectors have period matching the expected period of the solenoid as determined by LRRPredictor. Leading eigenvectors of graph Laplacian are periodic and are *π/*4-phase shifted, thereby yielding projections of LRR coil onto a winding around a circle in a 2D-plane. Phase estimation using the formula 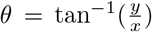 of LRR coil at bottom taking values between *−π* and *π*.

Generically, the leading pair of 0^th^ and 1^st^ eigenvectors of the graph Laplacian are out-of-phase periodic functions with frequency matching the expected frequency of the LRR coil (Figure 5), and thus yield projection onto principal axes perpendicular to the core of the coil (Figure 5). Given these *x* and *y* coordinates, we compute phase estimation *θ* to obtain a phase estimation along the LRR coil as shown in Figure 5 below.

## Results

### Cumulative winding number reveals errors made by ML-based LRR repeat unit delineator

We ran the LRR annotation tool LRRPredictor [13] on the 127 NLRs from *A. thaliana* to obtain predicted locations of the LRR motif “LxxLxL”. Let *R*_1_, …, *R*_*k*_ denote the starting residues for the LRR motifs predicted by LRRPredictor. The analogous measurement in our model is to record the residues at which our cumulative winding number *w* crosses integers.

To compare the two prediction schemes, we evaluate our cumulative winding number at the residues returned by LRRPredictor. That is, we form the list of numbers (*w*(*R*_1_), …, *w*(*R*_*k*_)). If the models are in agreement, the running difference (*w*(*R*_2_) *− w*(*R*_1_), …, *w*(*R*_*k*_) *− w*(*R*_*k−*1_) should equal the all-ones vector (1, …, 1) (that is, the structure should wind exactly once around the core between residues *R*_*j*_ and *R*_*j*+1_). The “discrepancy”

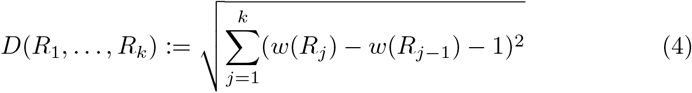

quantifies the extent to which this is not the case. A number of LRRPredictor outputs contained false predictions in which consecutive motif start sites *R*_*j*_ and *R*_*j−*1_ appear close together—often only a couple residues apart. Such duplicate predictions result in a high discrepancy *D*(*R*_1_, …, *R*_*k*_) because the difference *w*(*R*_*j*_) *− w*(*R*_*j−*1_) as computed in formula (4) above is close to 0.

To test the validity of our winding number computation, we ran the discrepancy computation on the LRRPredictor outputs on the 127 *A. thaliana* reference proteome NLRs as well as the training dataset for LRRpredictor, a manually-annotated “ground truth” dataset of LRR motifs on 172 experimentally-derived LRR structures taken from Protein Data Bank. These PDB protein structures were derived from a diverse set of organisms comprising bacteria, fungi, plants, and animals.

We found consistently low discrepancy values for the ground truth set with mean 0.127. By comparison, *A. thaliana* NLRome discrepancy values were generally low with mean 0.373, but exhibited higher values in cases where LRRpredictor made mistakes. Figure 6 below shows a pair of overlaid histograms comparing discrepancy values for both the validation dataset and NLRome dataset (S1 and S2 Table). The discrepancy values are much lower on the LRRPredictor ground truth dataset compared to the NLRome dataset, implying that our technique makes fewer mistakes than LRRPredictor does on new data. Figure 7 demonstrates how the discrepancy is able to catch duplicate motif predictions made by LRRPredictor. These results demonstrate not only the winding number’s ability to accurately model the LRR coil, but also its generalizability to non-NLR LRR’s derived from species other than *A. thaliana*.

**Figure 6:**
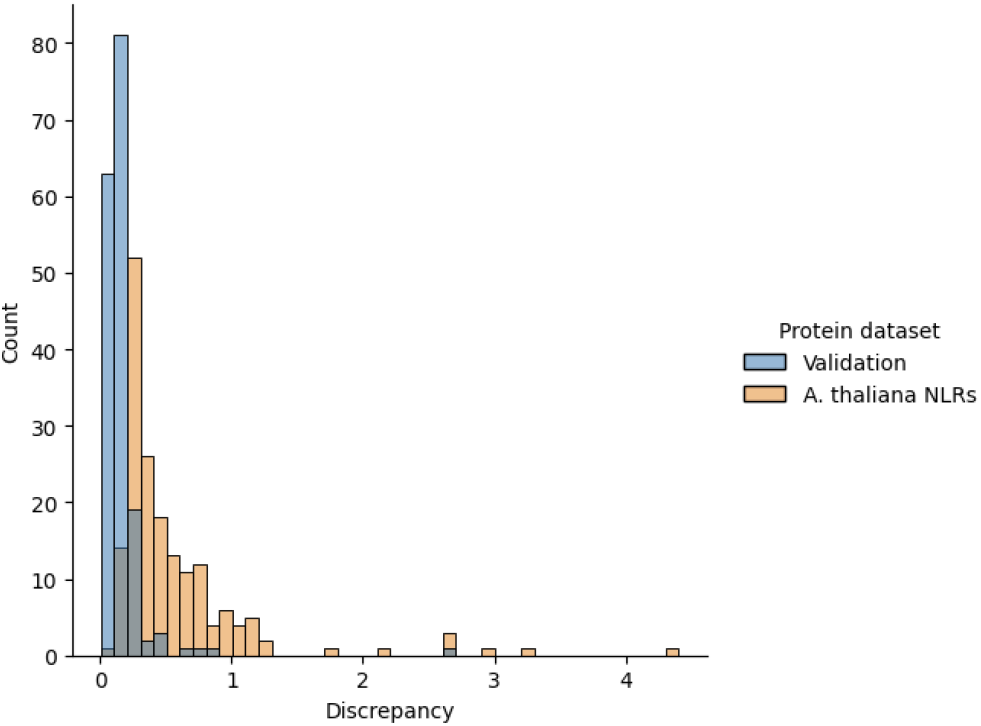
In yellow, histogram of discrepancies for LRRPredictor outputs on 127 *A. thaliana* NLRs showing a large peak around the mode. In blue, a histogram of discrepancies for manually-annotated LRR repeat units used as the training set for the LRRpredictor model. This ground truth dataset produces low overall discrepancy compared to LRRpredictor model outputs, thereby demonstrating the ability of the cumulative winding number computation to faithfully recapitulate the periodicity of the LRR coil.

**Figure 7:**
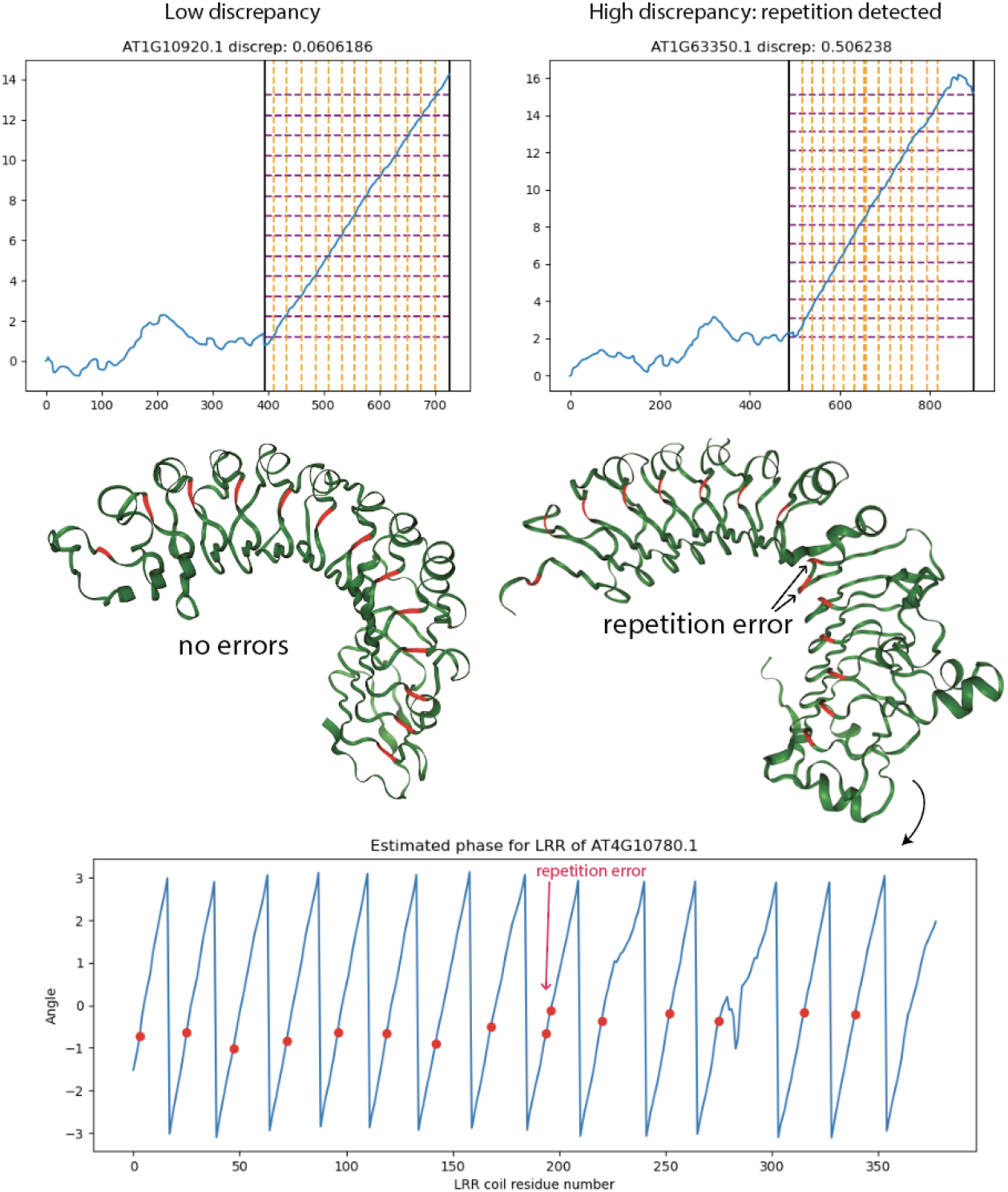
LRRPredictor discrepancy computation reveals proteins with erroneously repeated predictions. NLRs with high-discrepancy LR-RPredictor outputs tend to carry repetition errors or missing motif annotations. Orange vertical lines overlaid on winding number plot depict LRRPredictor residues, while purple horizontal lines depict the integer-spaced grid which best approximates the winding number graph evaluated at LRRPredictor residues. A repetition error can be seen in the grid representation as a doubled orange line around residue 685. At bottom, LRRPredictor residues are mapped onto graph Laplacian eigenvector phase estimation, revealing an pair of duplicates with adjacent phase.

### Structural anomaly detection by sliding window L2 distance from Laplacian eigenvector winding number to line

Many LRR coils have hairpin loops and other structural anomalies which deviate from coiling. In these anomalous regions, the leading eigenvectors deviate from their usual periodic behavior. Applying the winding number formula (2) above to the pair of leading graph Laplacian eigenvectors leads to a cumulative winding number within the LRR domain which is better able to discern small hairpins compared to the previous winding number computation based on normal bundle projection. As shown in Figure 8 below, we detect a small hairpin as a spike in L2 distance between the winding number and its median slope.

**Figure 8:**
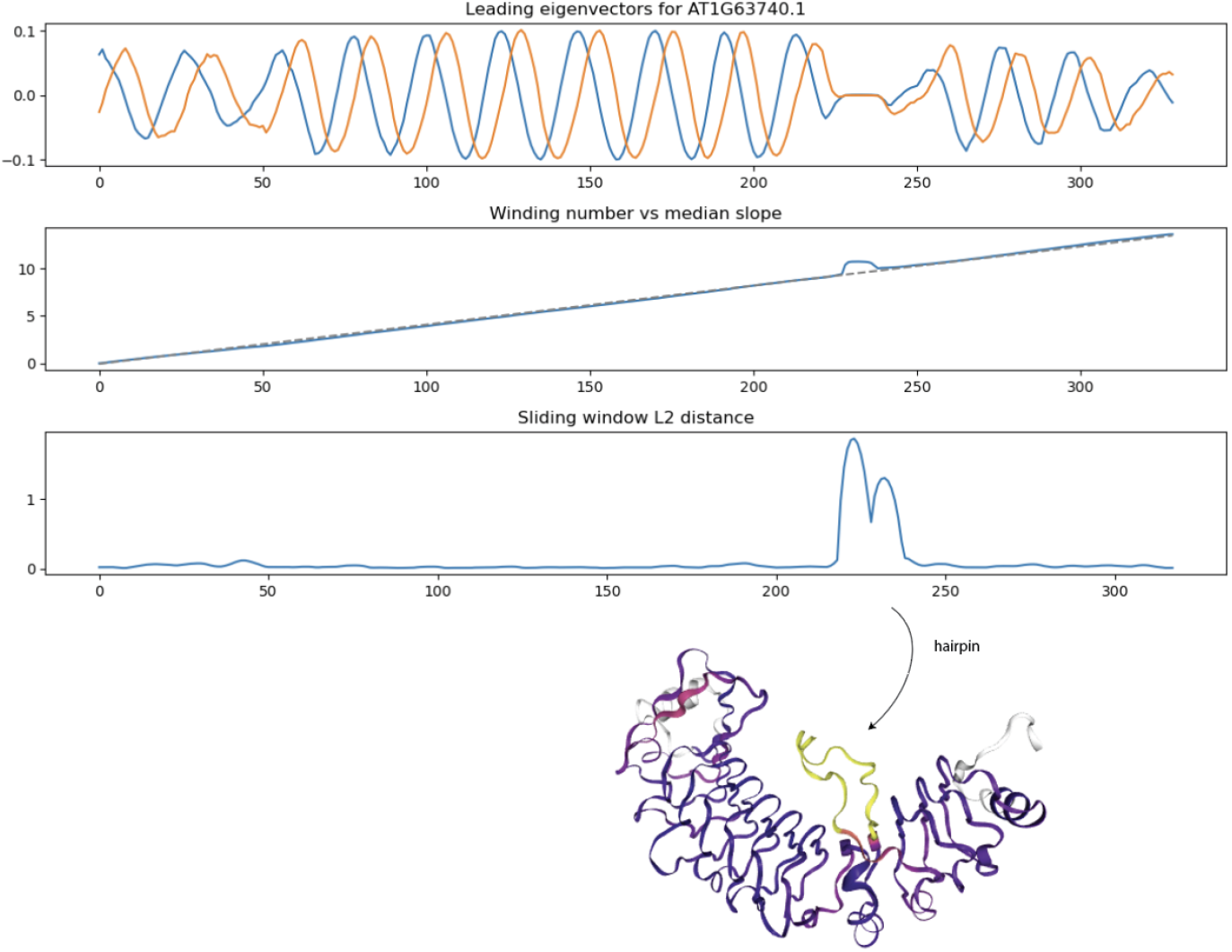
Sliding window L2 distance (SWL2D) from winding number to median secant line detects small hairpins/insertions in LRR coil domain. Structure at bottom is colored according to SWL2D where yellow values are higher.

## Discussion

The emergence of AlphaFold 2 has catalyzed a paradigm shift in protein structure prediction, facilitating access to genome-wide high-quality structural predictions.Traditional sequence homology-based domain annotation techniques, like LRRPredictor, often face challenges with LRRs, especially in proteins with high divergence and significant mutations. While evolutionary divergence might veil the sequence homology of LRR units, their core structural topology, characterized by 20-30 amino acid stretches typically involved in protein-protein interactions, often remains conserved, acting as a distinct structural signature. This study uses AlphaFold 2 to generate a 3D space curve from a protein sequence, which subsequently is projected into the 2D plane by identifying a series of “slinky” cross-sections. Through computing the cumulative winding number on the resultant 2D curve and employing piecewise linear regression, the linearly sloped region, identified as the LRR domain, is discerned. Our method pivots on the nuanced application of geometric data analysis to illuminate structural motifs that remain elusive to sequence analysis alone.

Our method yields several kinds of precise results: a) it identifies the start and end sites of the LRR domain with greater accuracy than HMM-based methods, b) it annotates repeat units more reliably than the existing LRRPredictor, c) it identifies misannotations by other annotation/prediction tools, and d) it reveals structural anomalies within the LRR domain that deviate from conventional coiling behaviors. These findings not only underscore the utility of our approach but also present a robust framework for delving into the intricate structural patterns intrinsic to LRR domains.

Our methods are general enough to adapt well to the detection of other solenoid domains. The outcomes from our approach serve as a foundation for structure-guided annotation of proteins containing LRR or other solenoid domains, which are often elusive to HMMs. In the broader context, our findings offer a comparative lens through which the evolution and function of NLRs, including repeat shuffling, can be scrutinized across various lineages [21]. Moreover, the precision and reliability of our annotation methodology can potentially serve as a catalyst for propelling research in disease resistance across a spectrum of plant species by furnishing detailed insights into the structure and function of LRR domains in NLR proteins.

Our method does come with limitations. For instance, while it can detect non-coiling structural anomalies within the LRR domain, the origin, authenticity, and potential functionality of these regions remain ambiguous. Moreover, our structure-based annotation method, albeit effective for domains with a straightforward geometric description like LRRs, might not be universally applicable to other protein domains without developing a new geometric model tailored to them. This underscores a potential limitation when juxtaposing sequence-based versus structure-based domain annotation, highlighting a future avenue warranting exploration: developing geometric models for other protein domains.

While we benchmarked our work on LRR domains in NLR proteins, the intrinsic methodology has the capacity for broader applications, extending to other solenoid protein domains like armadillo (ARM), tetratricopeptide (TPR), and ankyrin (ANK) repeats, all of which feature distinctive repeat sequences and structural configurations. The amalgamation of advanced protein structure prediction technologies and nuanced mathematical models, as demonstrated in our approach, underscores the potential for widening our understanding of protein function across varied biological systems.

## Declarations

## Acknowledgements

We thank Daniil Prigozhin for running LRRPredictor on *A. thaliana* NLRome, Chandler Sutherland and the Krasileva Lab for providing feedback and suggestions on this project and resulting manuscript.

## Funding

BX has been supported by Department of Energy Computational Sciences Graduate Fellowship grant number DE-SC0020347. KVK has been supported by funding from the Innovative Genomics Institute (https://innovativegenomics.org/), the Gordon and Betty Moore Foundation (https://www.moore.org/), grant number 8802, and the National Institute of Health New Innovator Director’s Award, grant number DP2AT011967. The funders had no role in study design, data collection and analysis, decision to publish, or preparation of the manuscript.

## Code availability

A Jupyter notebook for running the winding number LRR annotator is available at https://github.com/amcerbu/LRR-Annotation/tree/main

## Author contributions

BX contributed to conceptualization, data curation, formal analysis, methodology, investigation, software, validation, visualization, and writing. AC contributed to formal analysis, methodology, software, visualization, and writing. DL contributed to data curation, formal analysis, investigation, methodology, and validation. CJT contributed to formal analysis, methodology, software, visualization, validation, and writing. KK contributed to conceptualization, funding acquisition, supervision, and writing. BX wrote original draft of manuscript and other authors contributed to revisions.

## Notes

### Competing Interest Statement

The authors have declared no competing interest.

